# Recombinant SARS-CoV-2 spike S1-Fc fusion protein induced high levels of neutralizing responses in nonhuman primates

**DOI:** 10.1101/2020.04.21.052209

**Authors:** Wenlin Ren, Hunter Sun, George F. Gao, Jianxin Chen, Sean Sun, Rongqing Zhao, Guang Gao, Yalin Hu, Gan Zhao, Yuxin Chen, Xia Jin, Feng Fang, Jinggong Chen, Qi Wang, Sitao Gong, Wen Gao, Yufei Sun, Junchi Su, Ailiang He, Xin Cheng, Min Li, Chenxi Xia, Maohua Li, Le Sun

## Abstract

The COVID-19 outbreak has become a global pandemic responsible for over 2,000,000 confirmed cases and over 126,000 deaths worldwide. In this study, we examined the immunogenicity of CHO-expressed recombinant SARS-CoV-2 S1-Fc fusion protein in mice, rabbits, and monkeys as a potential candidate for a COVID-19 vaccine. We demonstrate that the S1-Fc fusion protein is extremely immunogenic, as evidenced by strong antibody titers observed by day 7. Strong virus neutralizing activity was observed on day 14 in rabbits immunized with the S1-Fc fusion protein using a pseudovirus neutralization assay. Most importantly, in less than 20 days and three injections of the S1-Fc fusion protein, two monkeys developed higher virus neutralizing titers than a recovered COVID-19 patient in a live SARS-CoV-2 infection assay. Our data strongly suggests that the CHO-expressed SARS-CoV-2 S1-Fc recombinant protein could be a strong candidate for vaccine development against COVID-19.

**Highlights:**

1. CHO-expressed S1-Fc protein is very immunogenic in various animals and can rapidly induce strong antibody production
2. S1-Fc protein solicits strong neutralizing activities against live virus
3. Stable CHO cell line expressing 50 mg/L of S1-Fc and a 3,000 L Bioreactor can produce 3 million doses of human COVID-19 vaccine every 10 days, making it an accessible and affordable option for worldwide vaccination

## Introduction

The SARS-CoV-2 was first identified in Wuhan, China at the end of 2019[1–5]. In four short months, the virus has caused a global pandemic, with over 2,000,000 confirmed cases and over 126,000 deaths worldwide. As a novel coronavirus with no effective treatments or drugs currently available, a vaccine is in dire need of development. Several broad approaches to the development of a COVID-19 vaccine have emerged, including DNA vaccines, RNA vaccines, viral vector vaccines, recombinant subunit vaccines, and dead viral preparations[6]. Among these, an RNA vaccine from Moderna was the first to reach human trials in early March in the US, followed by Cansino’s adenoviral vector vaccine which began human trials in China later in the same month.

Considering the spike protein is the receptor-binding protein that mediates viral-cell fusion during the initial infection event[7, 8], it has been favorably selected as a primary target for vaccine design. The spike protein has a total of 1,273 amino acids, which can be divided into two major domains according to their structures and functions[7, 9]. The first half is the S1 protein, which contains the RBD sequence and is located at the N-terminus of the spike protein[7, 10, 11]. The second half is the S2, serving as a trimeric structure that supports the S1 receptor binding site and has a fusion bundle which protrudes out into the host cell’s membrane after it is triggered by the S1 coming into contact with ACE II[9, 11, 12]. Due to the fact that most of the neutralizing epitopes are located within the S1 region, proteins containing the S1 region, the RBD, or even a trimeric approach to the S1+S2 have been considered as candidates for vaccine development[13].

In this study, we cloned the SARS-CoV-2 S1 protein (GenBank: QIC53204.1) as our vaccine candidate into a eukaryotic expressing construct and stably expressed CHO-K1 cells. The purified S1 protein was formulated with AD11.10 adjuvant before being immunized into mice, rabbits, and monkeys. Besides eliciting high levels of the anti-S1 antibodies, higher neutralizing activities against SARS-CoV-2 were also found in the anti-sera from monkeys when compared with the sera from a convalescent SARS-CoV-2 human patient. These results indicate that the S1 protein can effectively induce humoral immune responses in various animals and can elicit high levels of neutralizing antibodies in monkeys.

## MATERIALS AND METHODS

### Materials

AD11.10 (saponin based microemulsion) was from Advaccine Biopharma., China. Freund’s complete adjuvant (CFA) was purchased from SIGMA, USA. Female BALB/c mice were obtained from Vital River Co., China. New Zealand White rabbits were purchased and hosted at Longan Co., China. Cynomolgus monkeys were hosted at Xieerxin Biotech., China. Peroxidase conjugated secondary antibodies were sourced from Jackson Immunoresearch, USA. CHO-expressed SARS-CoV-2 S1-Fc fusion and S1-His6X were produced and purified by ZhenGe Biotech., China.

### Immunizations

4 weeks-old female Balb/c mice, 12-15 weeks-old female NZW rabbits and 3~4 years old Cynomolgus monkeys were immunized with CHO-expressed SARS-CoV-2 S1-Fc fusion after formulated with adjuvants according to manufacturer’s instructions. Blood samples were collected at different time points for measurement of antibody levels and neutralizing titers.

### Enzyme-linked immunoassay

The assay was carried out as described by Zhao RQ, et al[14]. Briefly, wells of 96-well plate were coated with 1.5 μg/mL of SARS-CoV-2 S1-His6X protein, blocked with 3% BSA-PBS. Serial dilutions of sera were first diluted with the normal sera from the same species, further diluted with Sample Dilution Buffer (20% Calf serum in PBS, 20%CS-PBS). 100μL of the diluted samples were transferred to each well of the plates. The captured antibodies were probed with HRP-conjugated secondary antibodies. After final washes, HRP substrate TMB solution (Beijing Kwinbon, China) was added and absorbencies were measured at 450nm with a microplate reader.

### Pseudovirus neutralization assay

HEK 293T cells and ACE2-transfected HEK 293T cells (ACE2-293T) were cultured in DMEM (Gibco, USA) supplemented with 10% heat-inactivated fetal bovine serum (FBS, Gibco), 50 U/ml penicillin (Gibco) and 50μg/ml streptomycin (Gibco) at 37°C with 5% CO2. HEK 293T cells were co-transfected with 10μg of a plasmid encoding SARS-CoV-2 S protein and 10μg of an env-deficient, luciferase-expressing HIV vector (pNL4–3.luc.RE) using Lipofectamine 2000 reagents (Life Technologies, USA). After 48 h, pseudovirus-containing culture supernatants were collected, filtered (0.45 μm), and stored at −80°C in 1ml aliquots.

ACE2-293T cells were seeded in 96-well plates at one day prior to infection. Heat-inactivated serum samples were 3-fold serially diluted and incubated with 200 TCID50 pseudoviruses for 1 h at 37°C. The mixtures were then used to infect ACE2-293T cells in triplicate. Following 12 h infection, wells were replenished with fresh medium and the luciferase activities of cells were determined 48 h later. Cells were lysed with lysis buffer (Glo-lysis buffer, Promega), followed by addition of luciferase substrate (Bright-Glo luciferase assay substrate, Promega). Luciferase activity was measured using a GloMax 96 microplate luminometer (Promega) and wells producing relative luminescence units (RLU) above three times of the mean background value were defined as positive. The 50% neutralization titer was calculated by probit analysis using the SPSS software.

### SARS-CoV-2 Neutralizing assay

Serum samples were inactivated at 56°C for 0.5h and serially diluted with cell culture medium in two-fold steps. The diluted serums were mixed with SARS-CoV-2 suspension of 100 TCID50 in 96-well plates at a ratio of 1:1, followed by 2 hours incubation at 36.5°C in a 5% CO2 incubator. 1-2 ×10^4^ Vero cells were then added to the serum-virus mixture, and the plates were incubated for 5 days at 36.5°C in a 5% CO2 incubator. Cytopathic effect (CPE) of each well was recorded under microscopes, and the neutralizing titer was calculated by the dilution number of 50% protective condition.

## RESULTS

### Production of SARS-CoV-2 S1-Fc fusion protein

The S1 protein (14Q-685R) was fused with Fc fragment of human IgG1 for ease of purification, and expressed by a stable CHO cell line. S1-Fc fusion protein was purified from the culture supernatant using a Protein-A column, as shown in Fig.S1A. The purity of S1-Fc protein was evaluated by running SDS-PAGE gel under reduced and non-reduced conditions, a band between 130-180 kD was detected as monomer of the S1-Fc, while the dimer appeared as a single band well above 180 kD molecule marker, suggesting the protein is heavily glycosylated (Fig.S1B).

### Immunizations of animals with SARS-CoV-2 S1-Fc fusion protein

The original purpose of this study was to generate positive control SARS-CoV-2 S1 antibodies for development of SARS-CoV-2 vaccine and serological assays. Strong adjuvants, such as Freund’s complete adjuvant (CFA) and AD11.10 (saponin-based microemulsion) were used to immunize animals.

Four 4-weeks old female Balb/c mice were first immunized with recombinant SARS-CoV-2 S1-Fc protein immersed in CFA and boost with S1-Fc protein in AD11.10 multiple times in short intervals. The tail bleeds were collected at different times and the antibody titers were examined by ELISA against purified SARS-CoV-2 S1-His6X. On day 7, all of the immunized mice showed syndrome of fevers, much earlier than usual, suggesting S1-Fc fusion protein is super immunogenic. The tail bleeds were collected right way and the anti-S1 antibody IgG titers were examined using HRP-conjugated goat anti-mouse IgG Fc-specific secondary antibodies. As shown in Fig. 1A, all of the four immunized mice already developed strong IgG titers (1:200 to 1:500) against SARS-CoV-2 S1 on Day 7. With one more boost on day 7, mouse #2’s anti-S1 IgG titer reached 1:5000 on day 12. It was sacrificed and the spleenocytes were used to generate mouse monoclonal antibodies. The remaining three mice also developed anti-S1 IgG titers higher than 1:10,000 on day 14 and day 20.

**Fig 1.**
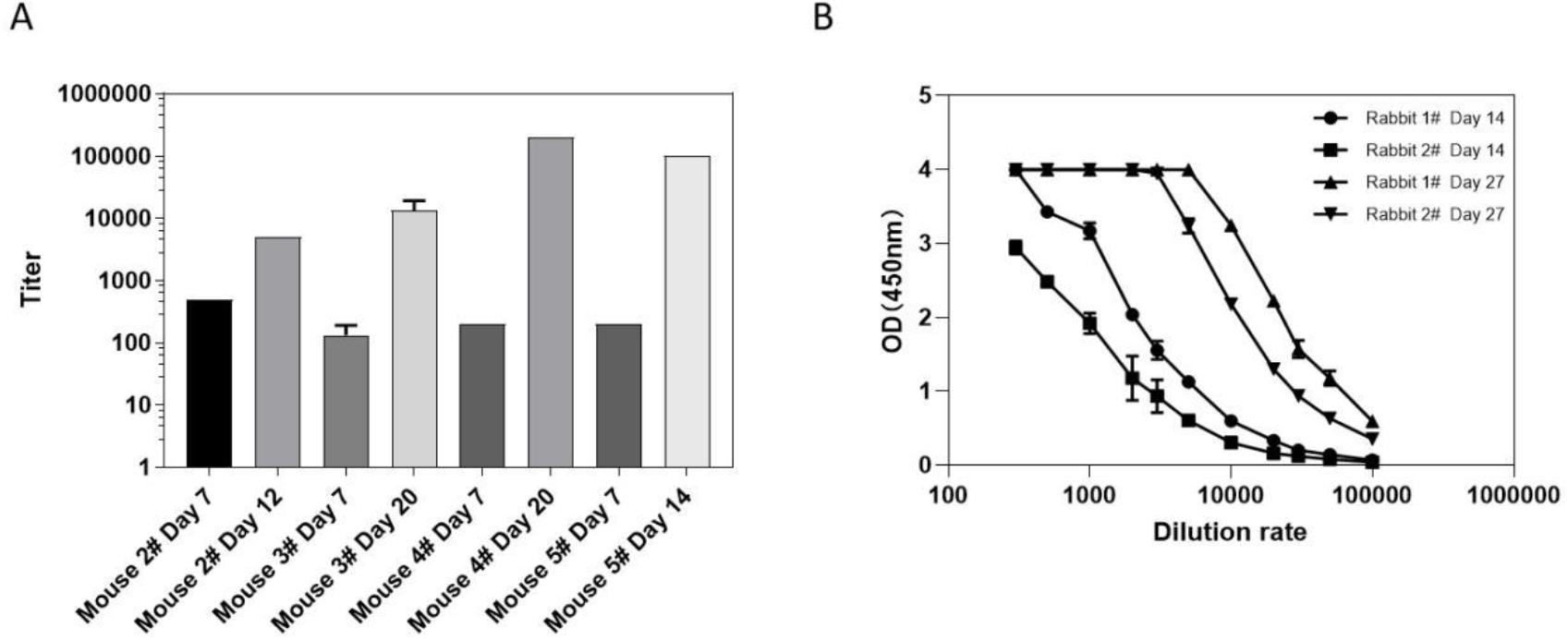
Anti-SARS-CoV-2 S1 antibody levels in S1-Fc immunized mice and rabbits. A) Serum samples collected from different mice on different days were evaluated with S1-His coated plates by ELISA using mouse IgG-specific secondary antibodies. The vertical coordinate is IgG titer. B) Serum samples collected from different rabbits on different days were evaluated with S1-His coated plates by ELISA using rabbit IgG-specific secondary antibodies. The horizontal coordinate is dilution and the vertical coordinate is absorbance value at 450nm.

The volume of mouse sera is too little to meet the needs for positive controls so two rabbits were immunized with S1-Fc fusion protein at a later day. On day 14 after multiple injections, similar to mice, the S1-Fc immunized rabbits already developed very high titers (1: 5,000 or higher) against the S1 protein (see Fig 1B).

In addition, one female and one male monkey were immunized with S1-Fc protein using CFA+AD11.10 (500μg each on Day 1, 5 and 23). The sera were collected on different days and evaluated by ELISA. HRP-conjugated goat anti-human IgG (H+L) secondary antibodies, which react with IgM, IgG1, IgG2, IgG4, were used for detection of total anti-S1 antibodies, while HRP-labeled anti-human IgG Fc and HRP-conjugated mouse anti-human μ-Chain were used for detections of IgG and IgM respectively.

As shown in Fig 2A, on day 9, anti-S1 total antibodies were detected only in the female monkey. On day 16, after two more boosts, good titers of anti-S1 total antibodies were observed in both monkeys. However, on day 23, there was a significant drop of total antibody titer in the female monkey. One boost was given on Day 24 and the total antibody titers in both monkeys increased greatly.

**Fig 2:**
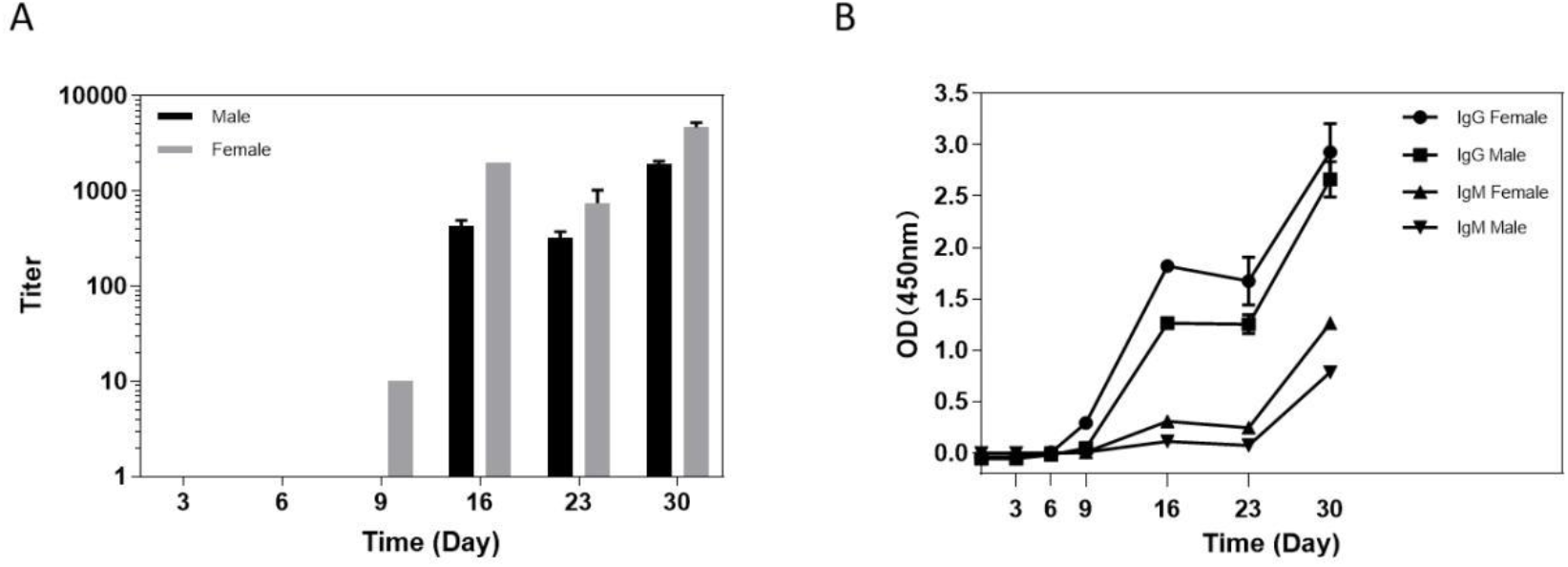
Anti-SARS-CoV-2 S1 antibody levels in S1-Fc immunized monkeys. A) The titers of total anti-S1 antibodies in monkey serum. Sera were evaluated by ELISA using HRP-conjugated goat-anti-human IgG (H+L) secondary antibodies. The horizontal coordinate is time and the vertical coordinate is the titer of anti-S1 antibodies in the sera. B) The levels of IgG and IgM in monkey sera. Sera were examined with either HRP-conjugated goat anti human IgG Fc-specific secondary antibodies (1:20000) or HRP-conjugated mouse monoclonal anti-μ-Chain of human IgM (1:5000). The horizontal coordinate is time and the vertical coordinate is absorbance value at 450nm.

When we used isotype-specific secondary antibodies to re-examine the sera, as shown in Fig. 2B, only anti-S1 IgG antibodies were seen in the female monkey on day 9. On day 16, strong anti-S1 IgG titers were detected in both monkeys, while the IgM levels were undetectable. Interestingly, after one more boost of S1-Fc on day 24, not only a big increase in anti-S1 IgG titers but also significant increase in anti-S1 IgM titers was observed. So when both S1-specific IgM and IgG were detected in the COVID-19 patients, it maybe be an indication either re-infection or re-emerging of SARS-CoV-2 from the infected human cells.

We also examined the levels of anti-S1 IgG and IgM antibodies in the plasma samples collected at different days post onset of COVID-19 disease from a recovered patient. As shown in Fig. 3, the IgG antibodies can be detected 4 days after onset and the titer increased on day 12, but reduced significantly on day 20 and 28. No IgM was detected.

**Fig 3.**
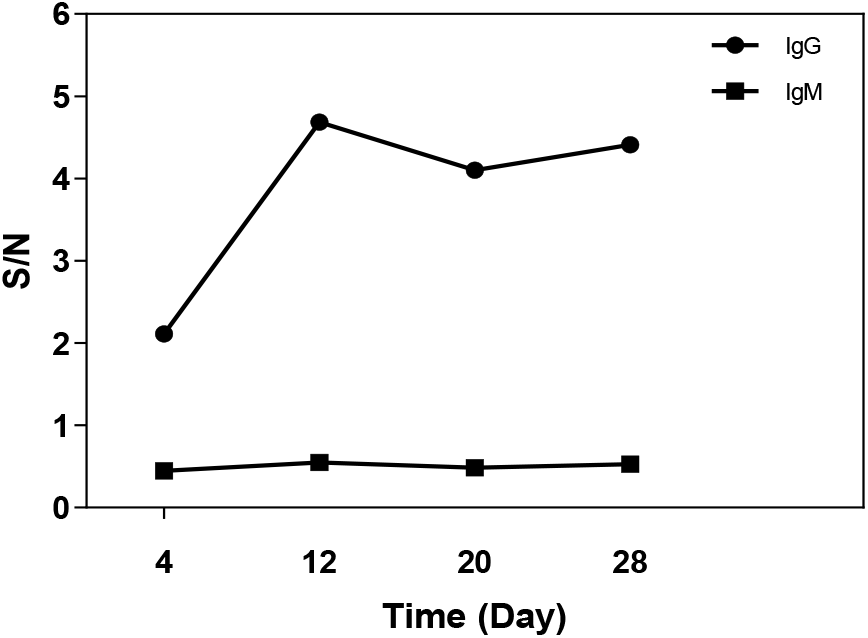
Levels of IgG and IgM in sera from COVID-19 convalescence patient. Sera were examined with either HRP-conjugated goat anti-human IgG Fc-specific secondary antibodies (1:20000) or HRP-conjugated mouse monoclonal anti-μ-Chain of human IgM (1:5000). The horizontal coordinate is time and the vertical coordinate is absorbance value at 450nm.

### Neutralization activities of SARS-CoV-2 S1-Fc fusion protein antisera

The virus neutralizing activities of the S1-Fc immunized sera were examined using either pseudo-virus or live SARS-CoV-2.

The neutralizing activity of the sera from S1-Fc immunized rabbits was examined using SARS-CoV-2 pseudo-virus. Briefly, heat-inactivated serum samples were 3-fold serially diluted and incubated with 200 TCID50 pseudo-viruses for 1 h at 37°C. The mixtures were then used to infect ACE2-293T cells. Three days later, the luciferase activities of the infected cells were measured. The 50% neutralization titer was calculated by probit analysis using the SPSS software. As shown in Fig. 4A, a neutralizing titer of 1:378 was obtained after the rabbits received three consecutive immunizations.

**Fig. 4.**
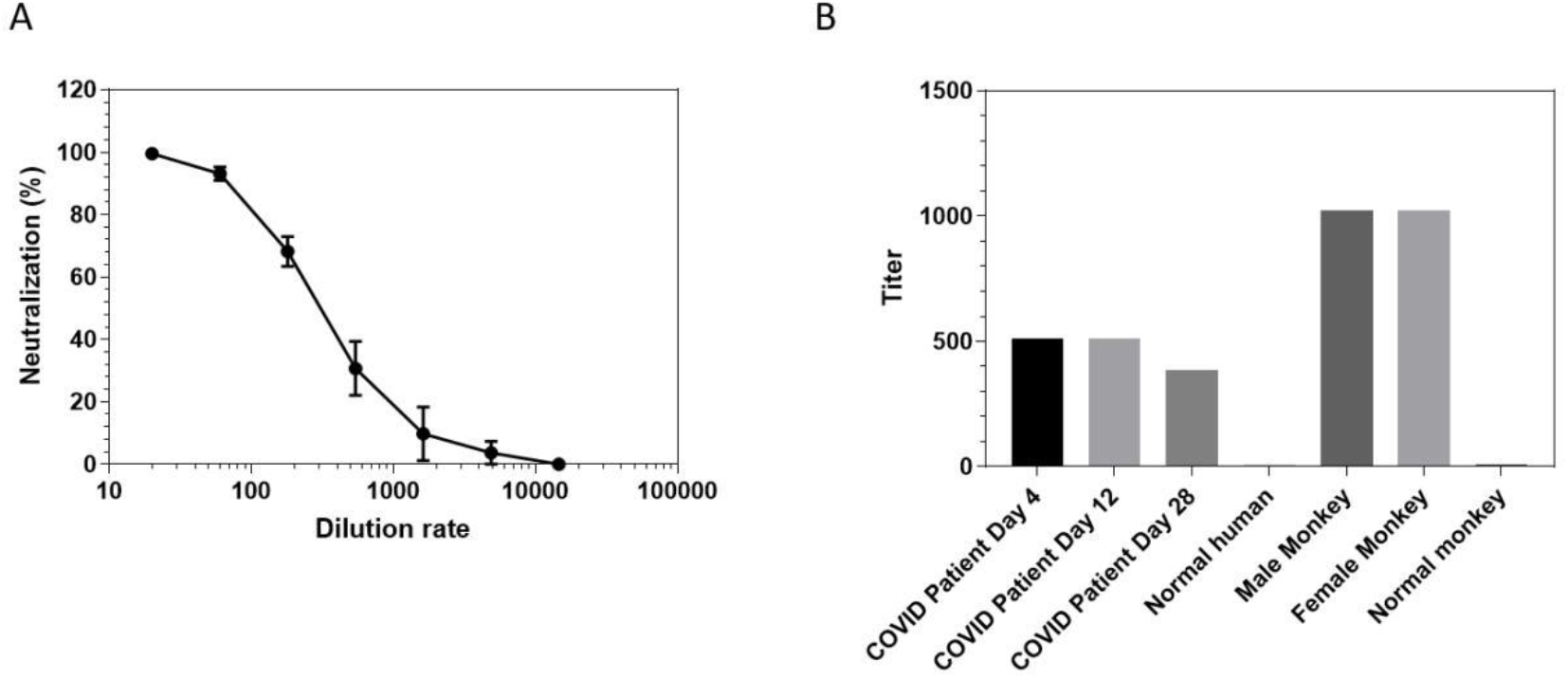
Virus neutralizing activities of antisera from either S1-Fc-immunized animals or COVID-19 convalescence patient. A) The neutralizing activity of rabbit sera was evaluated with SARS-CoV-2 pseudo-virus. The horizontal coordinate is Log10 serum dilution and the vertical coordinate is the percentage of the neutralization. B) The neutralizing titer of monkey and COVID-19 Convalescent patient’s sera was examined using live SARS-CoV-2. The vertical coordinate is the neutralizing titer.

We also compared the neutralizing titers of the sera from S1-Fc immunized monkeys with the ones of recovered COVID-19 patient’s plasma samples using live SARS-CoV-2. Heat-inactivated serum samples were first serially diluted with cell culture medium, pre-incubated live SARS-CoV-2 suspension of 100 TCID50 for 2 hours at 36.5°C. 1-2 ×10^4^ Vero cells were then added to the serum-virus mixture, and the plates were incubated for 5 days at 36.5°C. Cytopathic effect (CPE) of each well was recorded under microscopes, and the neutralizing titer was calculated by the dilution number of 50% protection. As shown in Fig. 4B, immunization of SARS-CoV-2 S1-Fc fusion protein induced very high neutralizing activities with titers >1:1024 in both monkeys against live SARS-CoV-2 infection, while the neutralizing titers of human sera from the recovered COVID-19 patient were at 1:512 on day 4 and day 12, but dropped to 1:384 on day 28 post the onset of disease, matched with the decrease of antibody levels in the plasma samples.

Our data clearly suggest the CHO-expressed full length SARS-CoV-2 S1-Fc fusion protein can solicit very strong neutralizing activities against SARS-CoV-2 in a very reasonable short of time and could be a good candidate for COVID-19 vaccine development.

## DISCUSSION

The spike protein of the SARS-CoV-2 has a total of 1,273 amino acids and two major domains according to their structures and functions. The first half is an S1 protein which contains the RBD sequence and is located at the N-terminus of the spike protein[7]. The second half is the S2, serving as a trimeric structure which supports the S1 receptor binding structure in place. Due to the RBD being located within the S1 region, we generated a S1-Fc fusion protein as a candidate for vaccine development.

Since the SARS-CoV-2 S1 protein is predicted to have at least 16 N-linked glycosylation sites[15], we chose mammalian cell CHO-K1 as the expression system to ensure the right glycosylation. CHO-K1 is also one of the most popular cell lines used to make human antibody drugs.

Shringrix, a vaccine based on the recombinant gE protein of varicella-zoster virus, was launched in 2018 by GSK for prevention and treatment of shingles with great success. The primary vaccination schedule consists of two dose of 50μg/0.5mL each two month apart[16]. Assuming the same vaccination schedule for COVID-19, with the stable expression level at 50mg/L of S1-Fc fusion protein, a 3,000 L CHO cell Bioreactor can easily produce 3 million dose of recombinant S1-Fc vaccine every two weeks. With so many antibody drug production plants in the world, it may be the ONLY feasible way to make enough COVID-19 vaccines for the entire world within one year.

There are many therapeutic biological drugs such as TNF Receptor-Fc fusion protein (Enbrel) that have been used for treatments of AMD and other human diseases without any known side-effect caused by the Fc part of the drugs. Dimerization of S1-Fc fusion protein via the disulfide bonds formed within the hinge regions of Fc may help to mimic the 3D conformation of oligomerized S protein in vivo. Most importantly, the human Fc domain fused to the C terminal of S1 protein greatly facilitates the purification of the recombinant protein following the standardized antibody drug manufacturing procedures.

Using the purified recombinant S1-Fc fusion protein as immunogen, we demonstrated it not only has very good immunogenicity in mice, rabbits and non-human primates, but also can elicit strong neutralizing activities against SARS-CoV-2 infection in vitro.

A study of its potential antibody-dependent enhancement (ADE) is currently underway. We have been using the SARS-S1-His6X protein to challenge the mice which have been immunized with S1-Fc fusion protein and exhibited strong anti-SARS-CoV-2 S1 titers. At this moment, we have not observed any abnormality with the mice.

### Conclusion

In this study, we established a CHO-K1 cell line that stably expresses the SARS-CoV-2 S1 protein with human IgG Fc fused to its C terminal. The purified recombinant S1-Fc fusion protein formulated with a saponin-based microemulsion adjuvant was used to immunize mice, rabbits and monkeys. Beside high levels of the anti-S1 antibodies elicited, higher neutralizing activities against live SARS-CoV-2 from the anti-sera of monkeys were developed within 16 days when compared with the sera from a recovered SARS-CoV-2 patient. Our results demonstrated that the S1-Fc fusion protein could effectively induce humoral immune responses in various animals and, significantly, can elicit high levels of neutralizing antibodies in non-human primates.

## Author contributions

WLR, RQZ, JXC, JGC, ALH, WG contributed the development of recombinant proteins. FF, STG, JCS,YLH, GZ, YXC,XJ, XC, ML, CXX contributed in the antibody generation and sample testing. GFG, BW, JX, GG helped in experiment designs. HS, SS, QW contributed in technical writing.

## Acknowledgments

This work is supported by Research Grants from Beijing Science and Technology Commission, and Bill & Melinda Gates Foundation to Le Sun. The funders had no role in the study design, data collection and analysis, decision to publish, or preparation of the manuscript. We thank Dr. Qiang Gao and his team from SinoVac for performing the live SARS-CoV-2 neutralizing assay, Dr. Jianqing Xu at Fudan Univ. for providing the ACE2-transfected HEK 293T cells, Dr. Pinliang Hu at Beijing Beyond Biotech. for purification of S1-Fc protein.

## REFERENCES

[1] Li Q, Guan X, Wu P, Wang X, Zhou L, Tong Y, et al. Early Transmission Dynamics in Wuhan, China, of Novel Coronavirus-Infected Pneumonia. The New England journal of medicine. 2020.

[2] Wu F, Zhao S, Yu B, Chen YM, Wang W, Song ZG, et al. A new coronavirus associated with human respiratory disease in China. Nature. 2020;579:265–9.

[3] Chan JF, Yuan S, Kok KH, To KK, Chu H, Yang J, et al. A familial cluster of pneumonia associated with the 2019 novel coronavirus indicating person-to-person transmission: a study of a family cluster. Lancet. 2020.

[4] Huang C, Wang Y, Li X, Ren L, Zhao J, Hu Y, et al. Clinical features of patients infected with 2019 novel coronavirus in Wuhan, China. Lancet. 2020.

[5] Li X, Zai J, Wang X, Li Y. Potential of large ‘first generation’ human-to-human transmission of 2019-nCoV. Journal of medical virology. 2020.

[6] Cohen J. Vaccine designers take first shots at COVID-19. Science. 2020;368:14–6.

[7] Li F. Structure, Function, and Evolution of Coronavirus Spike Proteins. Annual review of virology. 2016;3:237–61.

[8] Stertz S, Reichelt M, Spiegel M, Kuri T, Martinez-Sobrido L, Garcia-Sastre A, et al. The intracellular sites of early replication and budding of SARS-coronavirus. Virology. 2007;361:304–15.

[9] Wong SK, Li W, Moore MJ, Choe H, Farzan M. A 193-amino acid fragment of the SARS coronavirus S protein efficiently binds angiotensin-converting enzyme 2. The Journal of biological chemistry. 2004;279:3197–201.

[10] Ou X, Liu Y, Lei X, Li P, Mi D, Ren L, et al. Characterization of spike glycoprotein of SARS-CoV-2 on virus entry and its immune cross-reactivity with SARS-CoV. Nature communications. 2020;11:1620.

[11] Song W, Gui M, Wang X, Xiang Y. Cryo-EM structure of the SARS coronavirus spike glycoprotein in complex with its host cell receptor ACE2. PLoS pathogens. 2018;14:e1007236.

[12] Grant OC, Montgomery D, Ito K, Woods RJ. 3D Models of glycosylated SARS-CoV-2 spike protein suggest challenges and opportunities for vaccine development. bioRxiv. 2020:2020.04.07.030445.

[13] Ahmed SF, Quadeer AA, McKay MR. Preliminary identification of potential vaccine targets for the COVID-19 coronavirus (SARS-CoV-2) based on SARS-CoV immunological studies. bioRxiv. 2020:2020.02.03.933226.

[14] Zhao R, Li M, Song H, Chen J, Ren W, Feng Y, et al. Serological diagnostic kit of SARS-CoV-2 antibodies using CHO-expressed full-length SARS-CoV-2 S1 proteins. medRxiv. 2020:2020.03.26.20042184.

[15] Vankadari N, Wilce JA. Emerging WuHan (COVID-19) coronavirus: glycan shield and structure prediction of spike glycoprotein and its interaction with human CD26. Emerging microbes & infections. 2020;9:601–4.

[16] Label of SHINGRIX. 2017.

