## Supplemental files for "Recombinant SARS-CoV-2 spike S1-Fc fusion protein induced high levels of neutralizing responses in nonhuman primates"

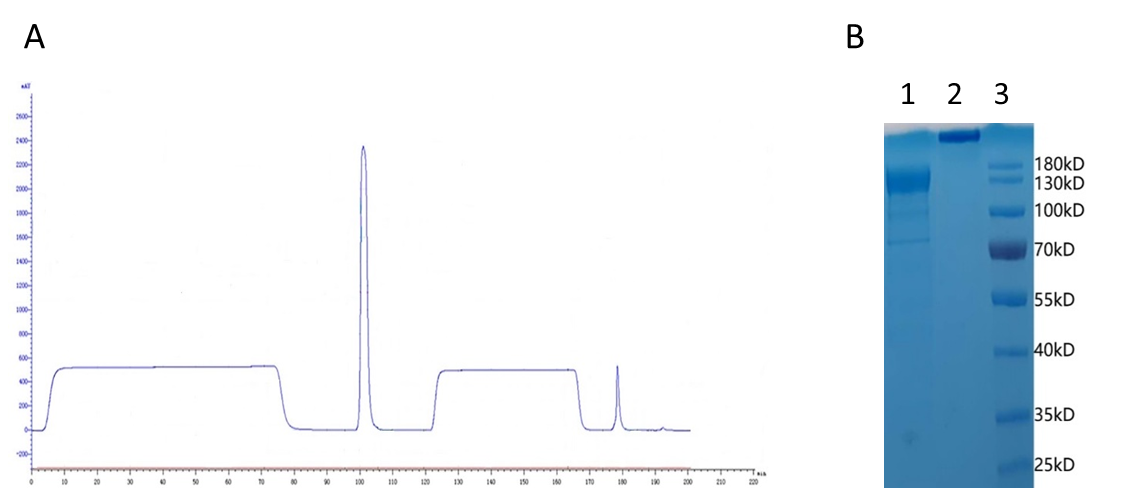

**Fig.S1 Purification of SARS-CoV-2 S1-Fc fusion protein.**

S1-Fc fusion protein was purified from the culture supernatant using Protein A column. A) The horizontal coordinate is time and the vertical coordinate is absorbance value at 280nm. B) SDS-PAGE analysis of purified S1-Fc fusion protein. Lane1, reduced S1-Fc protein; Lane 2, nonreduced S1-Fc; Lane 3 PageRuler™ Prestained Protein Ladder.
